# Selective *Shank3* Deletion in Glutamatergic Neurons of the Anterior Insular Cortex Induces Autism-Related Behavior and Circuit Dysfunction

**DOI:** 10.64898/2026.03.30.715416

**Authors:** Paula Mut-Arbona, Guilherme Horta, Zeina Msheik, Ignacio Marin-Blasco, Johans Pacheco-Villena, Tatiana Gusinskaia, Raul Andero, Luigi Bellocchio, Giovanni Marsicanno, Iñigo Ruiz de Azúa, Beat Lutz, Michael J. Schmeisser, Rafael Maldonado, Elena Martín-García

**Author notes:** These authors equally supervised this work (Rafael Maldonado, Elena Martín-García).

## Abstract

Mutations in the synaptic scaffold protein SHANK3 represent one of the most frequent genetic causes of autism spectrum disorder (ASD), yet the circuit mechanisms through which SHANK3 dysfunction leads to behavioral alterations remain incompletely understood. The anterior insular cortex (aINS) is a key integrative hub involved in socio-emotional processing, anxiety regulation, and social cognition, a group of behaviors frequently disrupted in ASD. Here, we investigated whether selective deletion of SHANK3 signaling in glutamatergic neurons of the aINS is sufficient to produce ASD-relevant behavioral and circuit phenotypes. Using conditional Shank3^flox4-22^ mice combined with stereotaxic viral delivery of Cre recombinase under the CaMKII promoter, we selectively deleted *Shank3* in glutamatergic neurons of the aINS. Behavioral phenotyping revealed increased anxiety-like behavior, enhanced repetitive behavior, and impaired social memory, while sociability and locomotor activity were largely preserved. These behavioral alterations were accompanied by genotype-dependent differences in neuronal activity revealed by calcium imaging, indicating disrupted activity dynamics in insular glutamatergic neurons following *Shank3* deletion. To assess the broader relevance of these findings, we evaluated the behavioral profile of BTBR T+ Itpr3tf/J mice, a model of idiopathic ASD, in the same battery of behavioral tests. Several behavioral alterations observed following insular *Shank3* deletion partially overlapped with those present in BTBR mice, supporting the relevance of aINS Shank3 in ASD-related phenotypes. Together, these findings identify glutamatergic neurons of the aINS as a critical locus through which Shank3 dysfunction can disrupt socio-emotional, cognitive, and repetitive behaviors. Our results highlight the aINS as a key circuit node contributing to ASD-related behavioral alterations and provide mechanistic insight into how synaptic scaffold disruption leads to circuit dysfunction and produces behavioral alterations.

## 1. Introduction

Autism spectrum disorder (ASD) is a highly multifactorial lifelong neurodevelopmental condition characterized by deficits in social interaction and communication, together with increased repetitive behaviors. In addition to these core symptoms, individuals with ASD frequently present with comorbid features including cognitive dysfunction, anxiety, sensory abnormalities, and motor impairments^1^. Despite substantial progress in identifying the complex polygenic architecture of ASD, the molecular and circuit mechanisms that link genetic alterations to behavioral phenotypes remain incompletely understood. Importantly, no pharmacological treatments currently approved by regulatory agencies effectively target the core symptoms of ASD. Indeed, existing therapies primarily address associated comorbidities such as hyperactivity, anxiety, and irritability, rather than the primary deficits of the disorder^2^.

Increasing evidence suggests that ASD involves disruptions in large-scale brain networks responsible for integrating emotional, sensory, and cognitive information^3,4^. Among these networks, the insular cortex has emerged as a key node in socio-emotional processing^5^. Notably, reduced connectivity within the insular network has been identified as one of the most consistent alterations across multiple genetically and idiopathically based mouse models of ASD^6^, highlighting the insular cortex as a convergent hub in ASD-related neurocircuitry. In particular, the anterior insular cortex (aINS) plays a central role in integrating interoceptive signals with emotional and motivational states through extensive connectivity with limbic structures including the anterior cingulate cortex and amygdala^7^. Consistent with this integrative function, neuroimaging studies have reported hypoactivation and disrupted functional connectivity of the aINS in individuals with ASD during tasks involving emotional and social processing^8,9^.

At the molecular level, alterations in synaptic scaffolding proteins have emerged as key contributors to ASD pathophysiology. Among these, the SHANK family represents one of the most prominent molecular substrates implicated in these disorders^10^. The *SHANK* family comprises three genes (*SHANK1, SHANK2*, and *SHANK3*), with *SHANK3* standing out as one of the strongest genetic risk factors for ASD^11^. Mutations in *SHANK3* occur in approximately 2% of individuals with ASD^12^, particularly in patients presenting with more severe clinical phenotypes, and lead to pronounced synaptic and behavioral abnormalities^13^. *SHANK3* encodes a multifunctional scaffolding protein located at the postsynaptic density of excitatory glutamatergic synapses, where it organizes large molecular complexes that regulate synapse formation, dendritic spine maturation, trafficking of glutamate receptors, and activity-dependent synaptic signaling^14^.

Genetic mouse models targeting *Shank3* have provided important insight into the neurobiological mechanisms underlying ASD. These models recapitulate core ASD-like phenotypes, including social deficits, repetitive behaviors, and anxiety-related alterations, as well as multiple synaptic abnormalities in excitatory circuits^15-17^. However, most studies to date have focused on global or regionally broad manipulations of SHANK3, leaving unresolved which specific neural circuits are sufficient to drive these behavioral phenotypes. Other mouse models of ASD have also been studied, such as the BTBR T^+^Itpr3^tf^/J mouse (BTBR)^18^. This highlights the polygenic origin of ASD but is consequently less appropriate to specific mechanistic studies.

Given the central role of the aINs in socioemotional integration and the importance of glutamatergic synaptic scaffolding in ASD, we hypothesized that disruption of Shank3 signaling in excitatory neurons of the aINS may represent a key circuit mechanism underlying ASD-related behavioral alterations. In the present study, we selectively deleted *Shank3* in glutamatergic neurons of the aINS using conditional genetic approaches combined with stereotaxic viral delivery of Cre recombinase. We then evaluated behavioral phenotypes relevant to ASD and assessed behavioral circuit-level consequences using calcium imaging on insular glutamatergic neurons. The behavioral results were compared to those obtained with the BTBR polygenic model of ASD. Our results identify glutamatergic neurons in the aINS as a critical locus through which Shank3 dysfunction could disrupt socioemotional behavior and circuit dynamics.

## 2. Materials and Methods

### 2.1. Animals

Shank3^flox ex4-22^ mice (Shank3 floxed exon 4–22, Jackson Strain # 032158, https://www.jax.org/strain/032158) were used to generate region-specific manipulations of *Shank3* expression^19^. To benchmark the behavioral phenotype against an idiopathic ASD model, BTBR T+ Itpr3tf/J mice and control C57BL/6J wild-type (WT) were also included.

Mice were housed in transparent Plexiglas cages under controlled environmental conditions (21 ± 1 °C; 55 ± 10% humidity) with food and water available *ad libitum*. Animals were maintained on a reversed 12 h light/dark cycle (lights off at 07:30; lights on at 19:30). All behavioral experiments were performed during the dark phase following a 30-min habituation period in the behavioral testing room.

All animal procedures were approved by the local ethical committee (Comitè Ètic d’Experimentació Animal – Parc de Recerca Biomèdica de Barcelona, CEEA-PRBB, 1649-(PG)EMG-22-0071-P2) and were conducted in accordance with ARRIVE guidelines and the European Directive 2010/63/EU for the protection of animals used for scientific purposes.

### 2.2. Viral vectors

Conditional Shank3^flox ex4-22^mice received stereotaxic injections of AAV-DIO-FLEX-GFP or AAV-CaMKII-Cre-mCherry. In animals injected with the latter AAV, Cre recombinase expressed under the CaMKII promoter selectively deletes Shank3 in glutamatergic neurons of the aINS. For calcium imaging experiments, Shank3 ^flox ex4-22^ mice were injected with an AAV expressing the calcium indicator GCaMP6f (AAV1/2-Syn-WPRE-SV40-GCaMP6f; Addgene, USA).

### 2.2. Experimental design

Male and female conditional Shank3 ^flox ex4-22^ and control-WT mice underwent stereotaxic viral injections targeting excitatory neurons in the aINS at postnatal day 24 to determine whether disruption of SHANK3 expressing glutamatergic neurons is sufficient to induce ASD-related phenotypes.

Two experimental groups of Shank3^flox ex4-22^ mice were generated (Fig. 1A): 1) Control-WT mice receiving a control AAV (AAV-DIO-FLEX-GFP) and 2) Glu-Shank3-KO-aINS mice in which *Shank3* was selectively deleted in glutamatergic neurons of the anterior insula via CaMKII-dependent Cre recombinase expression with an AAV-Cre (AAV-CamkII-Cre-mCherry). Following this strategy, Cre recombinase expressed under the CamKII promoter selectively deletes *Shank3* in glutamatergic neurons of the aINs.

**Figure 1.**
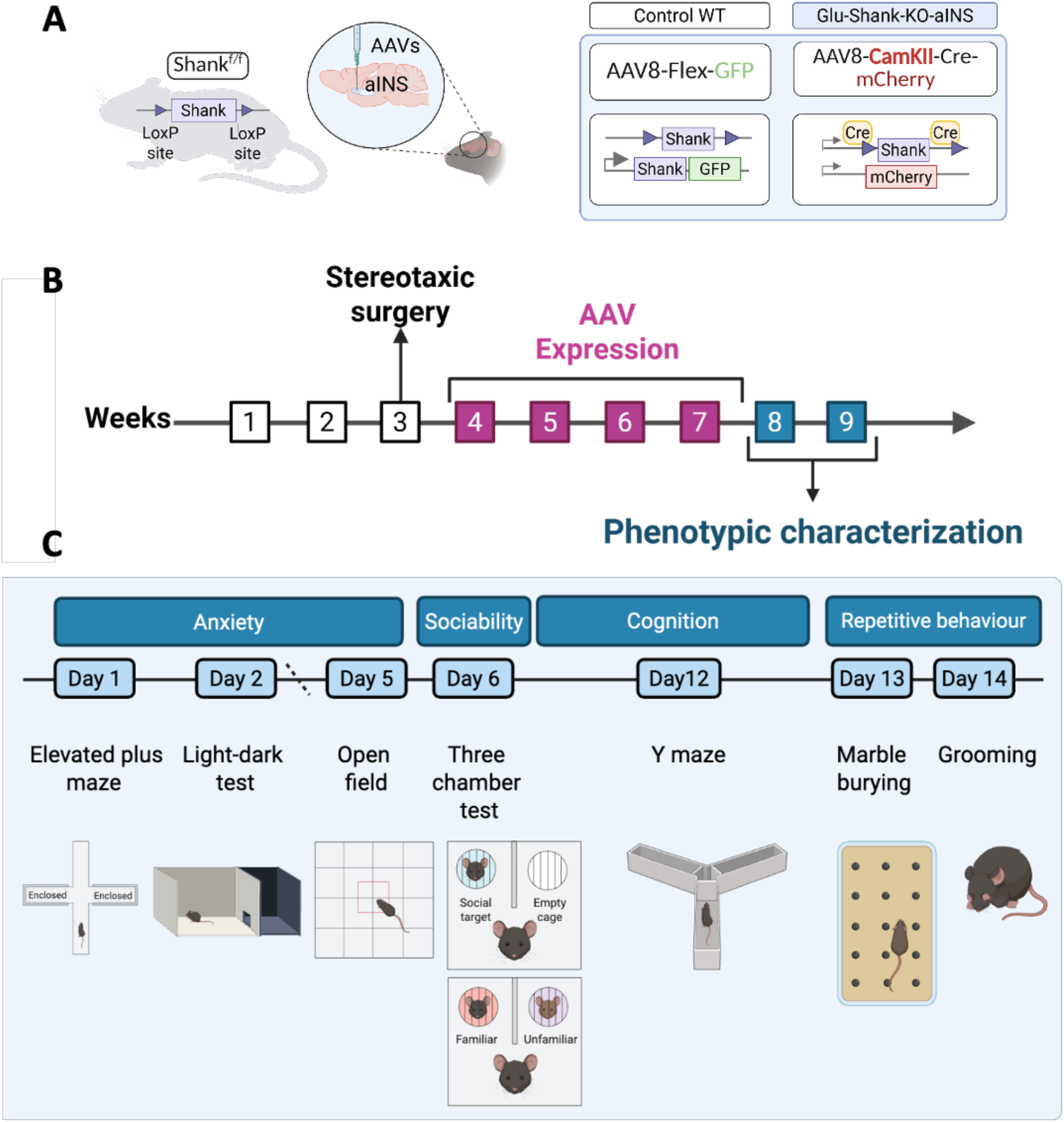
Cell-type-specific manipulation *Shank3* in glutamatergic neurons of the anterior insular cortex (aINS). (A) Schematic representation of viral vectors used for *Shank3* receptor deletion and subcellular rescue in glutamatergic neurons of the aINs for Contro-WT and Glu-Shank3-KO-aINs. (B) Overview of the behavioral testing pipeline used for phenotypic characterization following stereotaxic surgery and viral expression. Timeline of behavioral testing. Anxiety-like behavior was assessed using the elevated plus maze (day 1), light–dark box (day 2), and open-field test (day 5). Social behavior was evaluated using the three-chamber social interaction test (day 6). Cognitive performance was assessed using the Y-maze spontaneous alternation test (day 12). Repetitive behaviors were measured using the marble-burying test (day 13) and the grooming test (day 14). Behavioral sessions were recorded using the EthoVision XT 17 automated tracking system and analyzed blind to genotype and experimental condition.

A period of four weeks was allowed for viral expression before behavioral testing. Behavioral phenotyping began at approximately postnatal day 60.

Mice were behaviorally tested during the dark phase under controlled environmental conditions following a 30-min habituation period in the behavioral testing room. Phenotypic characterization was performed 4 weeks after surgery to allow for AAV expression (Fig. 1B). The behavioral pipeline was organized to minimize stress and carry-over effects, progressing from low-stress to more cognitively demanding tasks.

To evaluate the broader relevance of this circuit-specific model, BTBR mice and WT controls were subjected in parallel to the same behavioral testing battery as an idiopathic ASD comparison cohort.

Anxiety-like behavior was first assessed using the elevated plus maze (EPM), followed by the light–dark box and open-field test. Social behavior was evaluated using the three-chamber social interaction paradigm. Cognitive performance was assessed using the Y-maze spontaneous alternation test to evaluate working memory. Finally, repetitive behaviors were measured using marble burying and grooming tests (Fig. 1C).

All behavioral sessions were recorded using the EthoVision XT 17 automated video-tracking system and analyzed blind to genotypes and experimental conditions. At the end of the experiments (approximately 16 weeks of age), mice were deeply anesthetized and transcardially perfused for histological verification of viral targeting.

### 2.3. Stereotaxic surgery and viral vector microinjection

Stereotaxic surgeries were performed following the protocol previously described^20^ with minor modifications. Mice were anesthetized by intraperitoneal injection of ketamine hydrochloride (75 mg/kg) and medetomidine hydrochloride (1 mg/kg) diluted in sterile saline. Animals were placed in a stereotaxic frame (David Kopf Instruments, USA), and bilateral injections of adeno-associated viral vectors (AAV) were delivered into the aINS at postnatal day 23–25.

AAV gene delivery was performed using 33-gauge internal cannulas connected through polyethylene tubing to a 10 μl Hamilton microsyringe. A total volume of 0.2 μl per hemisphere was infused at a constant rate of 0.05 μl/min using a microinfusion pump. Following injection, the cannula remained in place for 10 min to allow diffusion and minimize reflux before slow withdrawal.

Injection coordinates targeting the aINS were adapted from the Paxinos and Franklin mouse brain atlas: anteroposterior (AP) +1.7 mm, mediolateral (ML) ±3.0 mm, relative to bregma, and dorsoventral (DV) −1.8 mm relative to brain’s surface. Postoperative care included subcutaneous administration of meloxicam (2 mg/kg) for analgesia and gentamicin (1 mg/kg) to prevent infection.

### 2.4. Behavioral testing

All behavioral experiments were conducted during the dark phase of the reversed light cycle following a 30-min habituation period in the testing room. Behavioral sessions were recorded using the EthoVision XT 17 automated tracking system and analyzed blind to genotype and experimental conditions.

#### 2.4.1. Elevated plus maze

Anxiety-like behavior was evaluated using the EPM. The apparatus consisted of four arms (29 × 5 cm) arranged in a cross configuration around a central platform (5 × 5 cm) elevated 40 cm above the floor. Two arms were open and 2 arms were enclosed by 20-cm-high walls. Light intensity was maintained at approximately 120 lux in the open arms and 5 lux in the closed arms. Mice were placed in the central square facing an open arm and allowed to explore freely for 10 min. The primary measures recorded were time spent in the open arms and the number of entries into the open arms. Mice that fell from the maze during testing were excluded from the analysis.

#### 2.4.2. Light–dark test

The light–dark test was used to evaluate anxiety-like behavior. The apparatus consisted of two compartments: a dark enclosed compartment and a brightly illuminated compartment (120 lux). Mice were placed in the dark compartment and allowed to habituate for 2 min with the door closed. The door was then opened, and mice were allowed to explore both compartments for 10 min. The time spent in each compartment was recorded.

#### 2.4.3. Open field

Locomotor activity was assessed in a white open-field arena (40 × 40 cm). Mice were placed in the center and allowed to explore for 10 min. Total distance travelled and time spent in the center were automatically recorded by the tracking system.

#### 2.4.4. Three-chamber social interaction test

Sociability and social novelty were assessed using the three-chamber test. The apparatus consisted of a Plexiglas box (50 × 25 × 30 cm) divided into 3 compartments connected by sliding doors. Cylindrical wire enclosures were placed in the center of each lateral chamber to contain the stimulus mice. The protocol consisted of three consecutive phases (10 min each). During the habituation phase, the test mouse was allowed to freely explore all three compartments. During the sociability phase, social interaction was assessed. A novel conspecific (Stranger 1), matched for sex and younger age, was placed under one wire enclosure, while the opposite enclosure contained a neutral object. In the social novelty phase, social memory was measured. A second unfamiliar mouse (Stranger 2) was placed in the previously empty enclosure while Stranger 1 remained in place. All stimulus mice were previously habituated to the enclosures to minimize stress and prevent active social engagement. The apparatus was not cleaned between phases to preserve relevant olfactory cues. Social behavior was quantified as time spent interacting with a stimulus mouse or object placed under wire enclosures.

#### 2.4.5. Y-maze spontaneous alternation

Spatial working memory was assessed using the Y-maze spontaneous alternation test. The maze consisted of three arms (arms A, B, and C), of dimensions 35 × 5 × 15 cm, arranged at 120°. Mice were allowed to freely explore the maze for 8 min, and spontaneous alternation percentage was calculated based on sequential arm entries. The percentage of spontaneous alternation was calculated as:

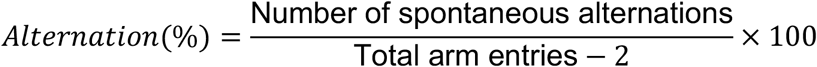

#### 2.4.6. Marble burying

Repetitive-like behavior was assessed using the marble burying test. Mice were placed in a transparent plastic cage (26.5 × 20 × 14 cm) containing bedding and 20 evenly spaced marbles and allowed to explore for 30 min. The number of marbles buried (defined as at least two-thirds covered with bedding) was counted by an experimenter blind to the experimental condition.

#### 2.4.7. Grooming

Spontaneous grooming behavior was assessed during a 10-min observation period in a clean cage. Grooming episodes were manually scored and categorized as head, face, or body grooming

### 2.5. Operant conditioning

Reward-driven behavior, impulsivity-like responding, and motivation were assessed using operant conditioning chambers (Med Associates, USA) equipped with 2 nose-poke ports, a pellet dispenser, and a house light^21^. Responses in the active port triggered delivery of a chocolate-flavored pellet (20 mg), whereas responses in the inactive port had no programmed consequences and were used as a control measure of nonspecific activity. Mice were first habituated to the operant chambers and the reward pellets. Training began under a fixed-ratio 1 (FR1) schedule, during which each active nose poke resulted in the delivery of 1 pellet. Animals were trained under FR1 for five consecutive daily sessions to establish stable responding. Sessions lasted 2 h or until the maximum number of rewards was reached. Following the acquisition, the reinforcement requirement was increased to a fixed-ratio 5 (FR5) schedule, in which 5 active responses were required to obtain a single pellet reward. FR5 sessions were also conducted for 2 h per day. During both FR1 and FR5 sessions, a time-out period (10s) followed reward delivery during which the cue light remained on, and additional responses were recorded but did not produce reward delivery. Responses during this time-out period were used as a measure of impulsivity-like behavior.

To assess motivational drive for the reward, animals were subsequently tested on day 11 under a progressive ratio schedule. In this schedule, the response requirement to obtain a reward progressively increased according to a predefined algorithm until the animal ceased responding. The breakpoint, defined as the highest ratio of completed responses required to obtain a reward, was used as an index of motivation.

The following behavioral measures were analyzed: (i) reinforcement, defined as the number of active responses leading to pellet delivery; (ii) impulsivity-like behavior, defined as responses emitted during the time-out period; and (iii) motivation, measured as the breakpoint obtained in the progressive ratio test.

All behavioral sessions were recorded automatically by the operant chamber software Med-PC-5.

### 2.6. Calcium imaging

#### 2.6.1. Viral injection for GCaMP6f expression

Animals were anesthetized with isoflurane (IsoFlo, Proima Ganadera SL, Spain) in oxygen at 4% for induction and 2.0–2.5% for maintenance, with an airflow of 1.25 L/min. The head was positioned in a stereotaxic frame (Kopf Model 962, Harvard-Panlab, Barcelona, Spain) using bregma and lambda as reference points. The skull was exposed and cleaned with saline (0.9% NaCl). Animals received 800 nL of AAV expressing the calcium indicator GCaMP6f (AAV1/2-Syn-WPRE-SV40-GCaMP6f) at the following coordinates: AP +1.8 mm; ML ±2.7 mm; DV −3.0 mm, relative to bregma. Injections were performed at a rate of 0.07 μL/min using a 5 μL Hamilton syringe (75RN model; Cibertec-Harvard, Madrid, Spain) coupled to an injection pump (KD Scientific, MA, USA). After injection, the syringe was slowly withdrawn to prevent reflux. The incision was sutured, and animals were allowed to recover under close supervision for one week. Postoperative care included subcutaneous meloxicam (2 mg/kg) for analgesia and gentamicin (1 mg/kg) to prevent infection.

#### 2.6.2. GRIN lens implantation and baseplate mounting

After one week of recovery, animals were anesthetized and placed in the stereotaxic frame. Following skull exposure, a 0.5 mm drill bit was used to perforate the bone at the previously defined AP and ML coordinates. A guide tract was then created using a 0.5 × 16 mm blunt syringe needle to a depth of 100 μm above the viral injection site (DV −2.9 mm). A GRIN lens (1.0 pitch, 0.5 mm diameter, 4 mm length; Inscopix, CA, USA) was then implanted at DV −3.0 mm. Saline was continuously applied to the entry site to facilitate insertion. The lens was secured with cyanoacrylate adhesive, and two skull screws were placed as anchoring system. The exposed area was covered with dental cement (Rebaron, GC Dental) mixed with black ink (Edding T-100). The lens was protected with a plastic cap (0.2 mL PCR tube cap) fixed with cyanoacrylate.

Postoperative care included Meloxicam (2 mg/kg) and gentamicin (1 mg/kg). Animals were individually housed and monitored for 2 weeks. After this period, the cap was removed and a metallic baseplate was fixed using dental cement. During this procedure, calcium activity was monitored with the Miniscope to optimize focal plane alignment. Animals were allowed to recover for an additional week before behavioral testing.

After behavioral testing, mice were deeply anesthetized and transcardially perfused for histological verification of viral targeting. Representative image of Cre- and GCaMP6f-expressing neurons in the aINS is shown in Supplementary Figure 1.

#### 2.6.3. Calcium imaging recording and data analysis

Neuronal calcium transients were recorded during behavioral tests (open field, three-chamber social interaction, and marble burying) using a Miniscope (v4.0; UCLA, CA, USA) coupled to Miniscope DAQ hardware and acquisition software. Mice were habituated to the Miniscope by allowing free exploration of their home cage without food or water for 10 minutes per day over three consecutive days. Imaging parameters (LED power and gain) were adjusted to avoid signal saturation and kept constant across sessions.

Locomotion was monitored using the center of mass (COM) of each mouse as a positional reference. Behavioral states were identified using a Gaussian Hidden Markov Model (HMM) based on kinematic features extracted from the COM trajectory. This probabilistic model represents behavior as transitions between hidden states, where each state generates observations according to a multivariate Gaussian distribution defined by a mean vector and covariance matrix. The most likely sequence of states was inferred using the Viterbi algorithm. Based on locomotion dynamics, the HMM identified three behavioral states: rest, exploration, and active locomotion.

Calcium imaging data were processed to extract neuronal fluorescence traces and converted to ΔF/F_0_ signals. Calcium events were detected and converted into a binary event matrix representing the temporal occurrence of Ca^2+^ transients. The association between neuronal calcium events and locomotion states was quantified using the Phi coefficient (φ). φ values were converted into Z-scores, and neurons with φ values greater than +1.96 standard deviations were classified as active, whereas neurons with φ values below −1.96 standard deviations were classified as inactive. Neurons not meeting either criterion were classified as unresponsive.

For each locomotion state, the following parameters were calculated: mean ΔF/F_0_ fluorescence amplitude, area under the curve (AUC) of calcium transients, calcium event rate (events/s), and total number of calcium events. These metrics were compared across locomotion states and between experimental groups to evaluate genotype-dependent differences in neuronal activity.

### 2.7. Statistical analysis

Statistical analyses were performed using GraphPad Prism (version 11). Data distribution was assessed using the Shapiro-Wilk normality test to determine whether parametric or non-parametric tests were applied.

Comparisons involving two groups and time were analyzed using repeated measures two-way analysis of variance (ANOVA) followed by Newman-Keuls post hoc tests when appropriate. Comparisons between two groups were performed using an unpaired Student’s t-test (parametric) or U-Mann-Whitney (non-parametric). The chi-squared test was used to compare differences in percentages between groups. Data are presented as individual values with mean ± SEM. Statistical significance was defined as p < 0.05.

## 3. Results

### 3.1. Selective *Shank3* deletion in insular glutamatergic neurons recapitulates core ASD-related behavioral alterations

Behavioral characterization revealed that deletion of *Shank3* in glutamatergic neurons of the aINS resulted in increased anxiety-like behavior. First, no significant differences were detected in the EPM, either in time spent in the open arms or in the number of open-arm entries (Fig. 2A–B). In contrast, anxiety-like phenotype was reflected by a significant reduction in the time spent in the light compartment of the light–dark test in Glu-Shank3-KO-aINS mice compared with Control-WT animals (p<0.01, Fig. 2C), suggesting that anxiety-related alterations were context dependent. Locomotor activity assessed in the open-field test did not differ between groups (Fig. 2D), indicating that general exploratory behavior and baseline locomotion were not affected by *Shank3* deletion.

**Figure 2.**
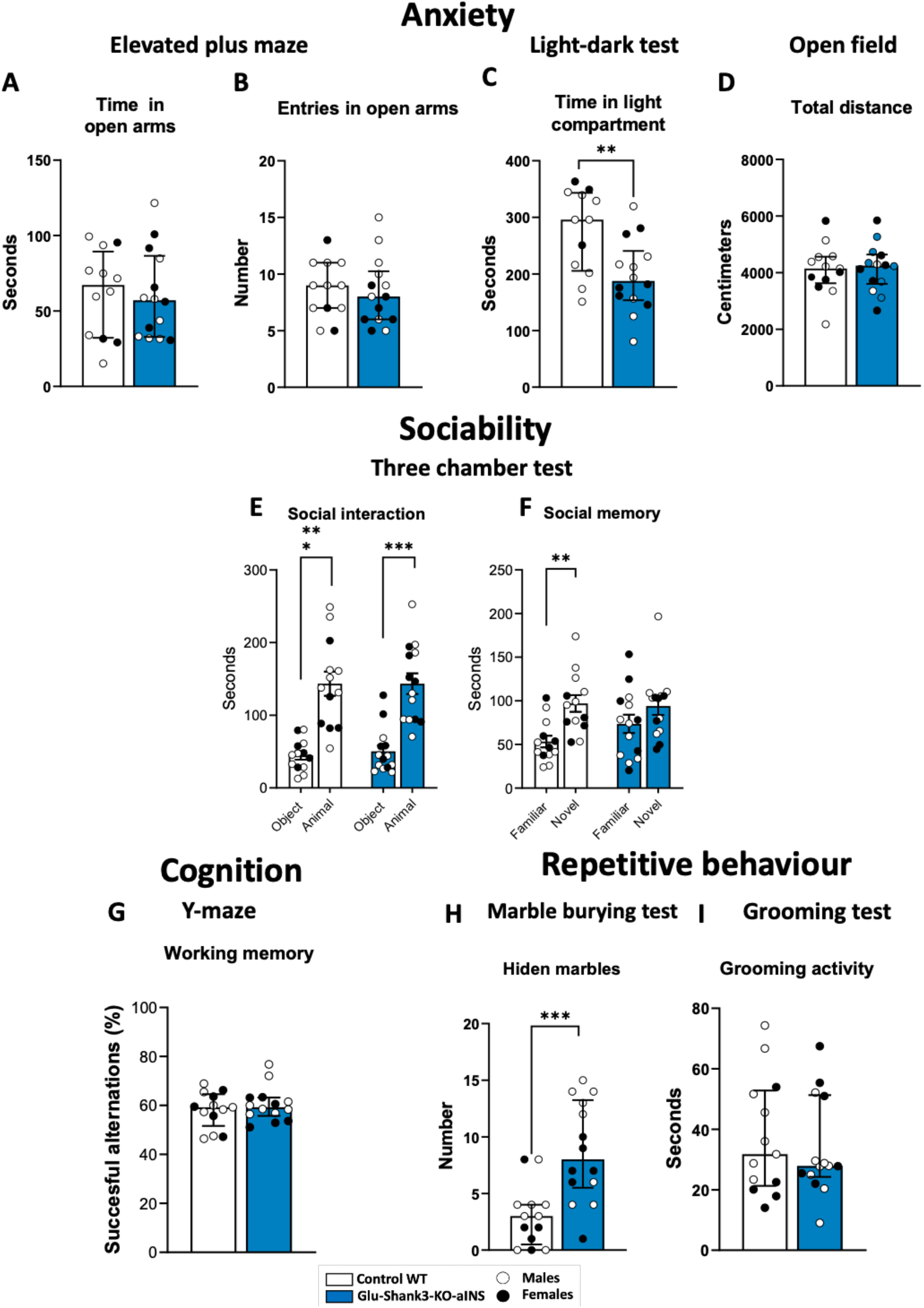
Anxiety, sociability, cognition, and repetitive behaviors following *Shank3* deletion in anterior insula glutamatergic neurons. Behavioral comparison was assessed between Glu-Shank3-KO-aINS and Control-WT mice. Anxiety-related behavior assessed via: (A) Time spent in open arms and (B) Number of entries into open arms in the EPM. (C) Time spent in the light compartment in the Light-Dark Box test and Total distance traveled in the Open Field test (locomotion). Social behavior assessed using the Three-Chamber Test (E) Social interaction and (F) Social memory. Cognition assessed via: (G) Working memory performance in the Y-maze. Repetitive behavior evaluated by: (H) Number of marbles buried in the Marble Burying test and (I) Duration of grooming. Data are expressed as mean + SEM (n= 13 for Control-WT and n= 14 for Glu-Shank3-KO-aINs). Data were analyzed using Student’s t-test or the Mann–Whitney U test, according to normality as determined by the Shapiro–Wilk test. Statistical significance was defined as **p < 0.01, ***p < 0.001.

Sociability, measured using the three-chamber social preference test, showed a similar differentiation between object and mouse in both groups, with a preference towards the mouse than the object marked by higher sniffing time versus an object (Fig. 2E). In contrast, social memory, assessed as the preference for a novel versus a familiar conspecific, was impaired in Glu-Shank3-KO-aINS mice compared with Control-WT animals as no statistically significant difference in sniffing time was observed between the novel and the familiar mouse (Fig. 2F), revealing a selective deficit in social recognition.

Working memory was evaluated using the Y-maze spontaneous alternation test. No significant differences were detected between Glu-Shank3-KO-aINS and Control-WT groups (Fig. 2G), indicating preserved working memory. In contrast, repetitive-like behavior was increased in Glu-Shank3-KO-aINS mice, as indicated by a higher number of buried marbles in the marble burying test (p<0.001, Fig. 2H), while grooming behavior remained unchanged (Fig. 2I).

Together, these findings demonstrate that selective deletion of *Shank3* in glutamatergic neurons of the aINs is sufficient to induce key ASD-relevant behavioral alterations, including increased anxiety-like behavior, impaired social memory, and enhanced repetitive behavior.

### 3.2. BTBR mice exhibit selective anxiety-related alterations without broad ASD-like phenotypes

To contextualize the behavioral effects observed in our brain-specific manipulations, we included the BTBR T+ Itpr3tf/J (BTBR) mouse line, an established idiopathic model of ASD, as a reference strain for phenotypic comparison. Consistent with previous reports, BTBR mice exhibited increased anxiety-like behavior, as indicated by reduced time spent in the light compartment of the light–dark test (p<0.05, Fig. 3C). However, no differences were observed in the EPM, either in time spent in open arms or in open-arm entries (Fig. 3A–B).

**Figure 3.**
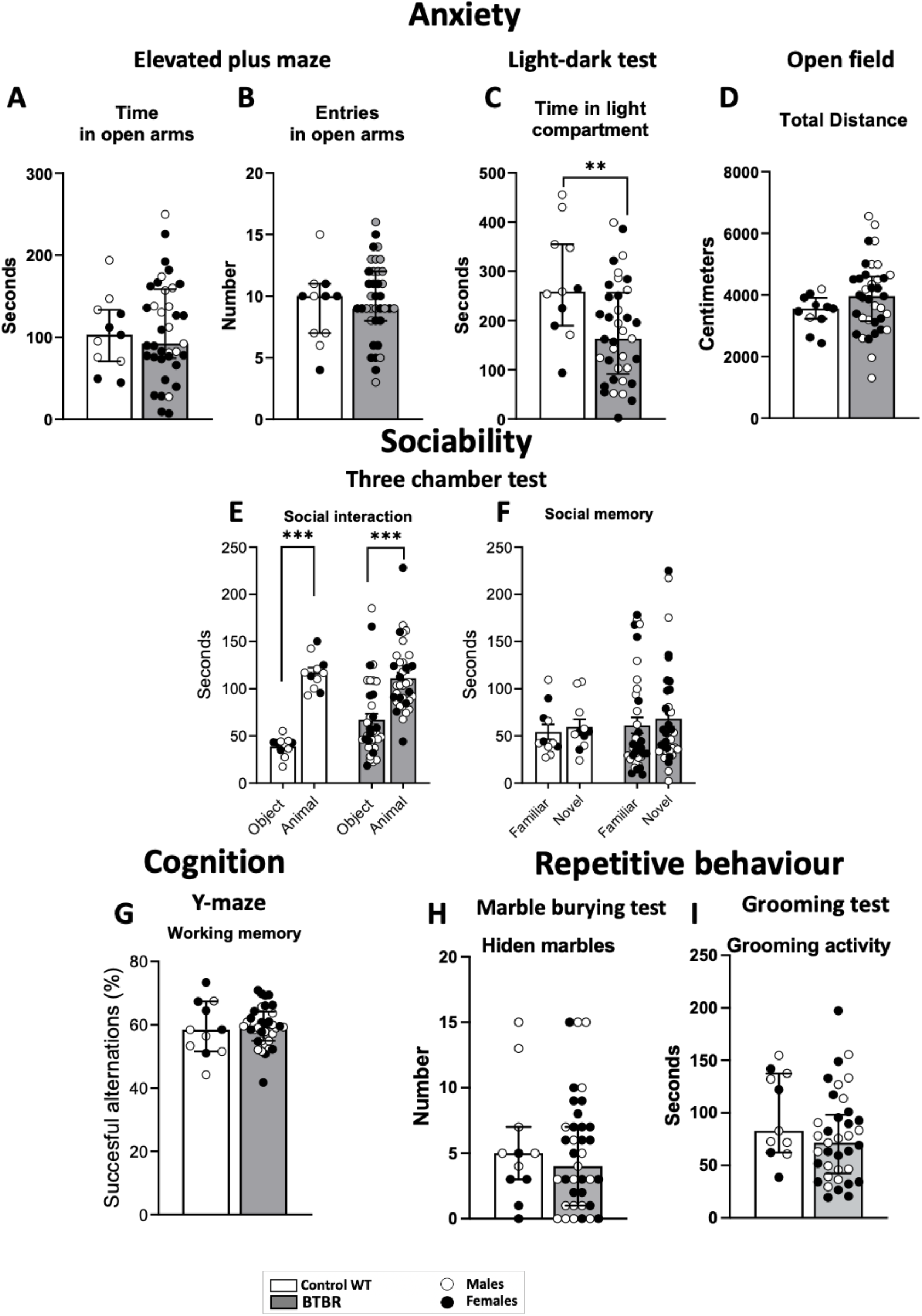
Anxiety, sociability, cognition, and repetitive behaviors in BTBR idiopathic model of autism. Anxiety-related behavior assessed via: (A) Time spent in open arms and (B) Number of entries into open arms in the EPM. (C) Time spent in the light compartment in the Light-Dark Box test and (D) Total distance traveled in the Open Field test (locomotion). Social behavior assessed using the Three-Chamber Test (E) Social interaction and (F) Social memory. Cognition assessed via: (G) Working memory performance in the Y-maze. Repetitive behavior evaluated by: (I) Number of marbles buried in the Marble Burying test and (J) Duration of grooming. Data are expressed as mean + SEM (n= 11 for Control-WT and n= 37 for BTBR). Data were analyzed using Student’s t-test or the Mann–Whitney U test, according to normality as determined by the Shapiro–Wilk test. Statistical significance was defined as **p < 0.01, ***p < 0.001.

Locomotor activity measured in the open-field test was comparable between BTBR and WT mice (Fig. 3D), indicating preserved baseline exploratory behavior. Both genotypes displayed similar social interaction in the three-chamber test marked by significantly higher sniffing time of the mouse rather than the object (Fig. 3E). However, social memory was not significantly altered in both groups under the conditions tested (Fig. 3F). Spatial working memory evaluated using the Y-maze spontaneous alternation test was also comparable between BTBR and WT animals (Fig. 3G). Similarly, repetitive behaviors measured through marble burying and grooming tests did not differ significantly between groups (Fig. 3H–I).

Together, these findings indicate that BTBR mice display anxiety-related alterations, but do not exhibit robust impairments across all behavioral domains examined in the present study, highlighting the selective nature of the phenotypes observed in our circuit-specific model.

### 3.3. Operant behavior is not altered by *Shank3* deletion in the anterior insular cortex

Reward-driven and impulsive behaviors were evaluated using operant conditioning paradigms (Fig. 4A–B). Significant differences were observed between Glu-Shank3-KO-aINS and Control-WT mice on day 3 (p<0.05) of FR1 reinforcement responding, with Glu-Shank3-KO-aINS showing lower levels of the primary reinforcing effects of chocolate-flavored pellets (Fig. 4C). Impulsivity-like behavior during the time-out period was also reduced in Glu-Shank3-KO-aINS with significantly lower levels in sessions 8 and 9 of FR5 (p<0.05, Fig. 4D). Motivation assessed using the progressive ratio test showed no significant differences between genotypes (Fig. 4E). Next, operant behavioral measures of reinforcement and impulsivity showed lower levels of primary reinforcing effects of chocolate-flavored pellets in session 5 (p<0.05) of FR1 in BTBR compared to WT mice (Fig. 4F) and lower impulsivity of these mutants in sessions 2-3 (p<0.05) and 5 (p<0.01) of FR1 (Fig. 4G) compared to WT mice. Importantly, BTBR mice showed lower motivation for chocolate-flavored pellets than control WT mice (p<0.05).

**Figure 4.**
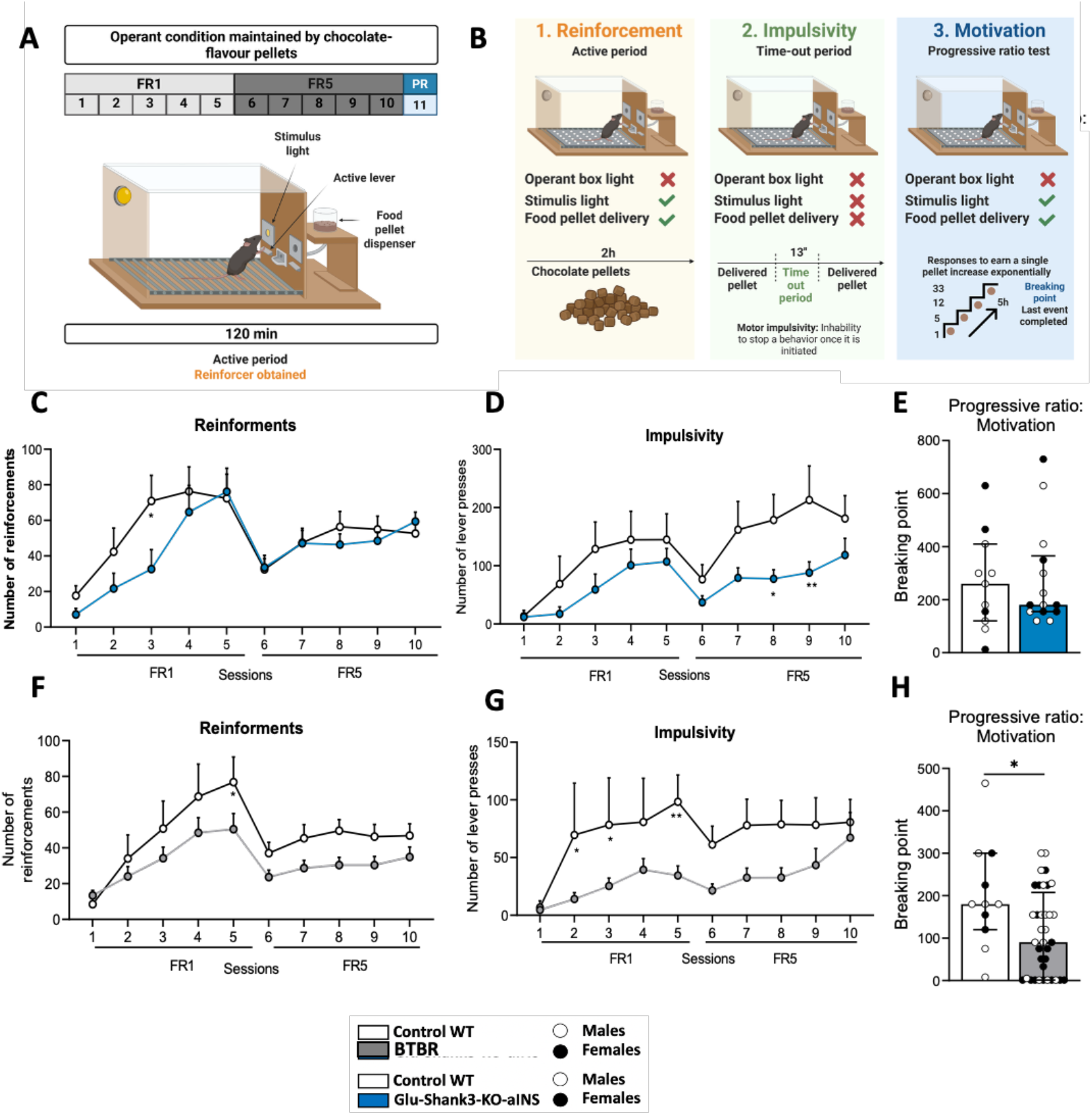
Operant conditioning assessment of reward-driven behavior, impulsivity, and motivation following *Shank3* deletion in anterior insular glutamatergic neurons and in the BTBR idiopathic model of autism. (A) Experimental schedule for operant conditioning maintained by chocolate-flavored pellets. Mice were trained under a fixed-ratio 1 (FR1) schedule for 5 days, followed by 5-days fixed-ratio 5 (FR5) sessions. On day 11, motivation was evaluated using a progressive ratio schedule. (B) Behavioral measures obtained in operant chambers, including reinforcement responding (number of active responses leading to pellet delivery), impulsivity-like responding during the time-out period, and motivation measured as breakpoint in the progressive ratio test. (C–E) Operant behavioral performance in Glu-Shank3-KO-aINS and Control-WT mice showing reinforcement responding (C), impulsivity-like responding (D), and motivation (E). (F–H) Operant behavioral performance in BTBR and WT mice showing reinforcement responding (F), impulsivity-like responding (G), and motivation (H). Data were analyzed using two-way repeated-measures ANOVA followed by Newman-Keuls post-hoc test. Student’s t-test or the Mann–Whitney U test was used when required, depending on normality as determined by the Shapiro–Wilk test. Data are expressed as mean + SEM. Statistical significance was defined as *p < 0.05.

These results indicate that selective deletion of *Shank3* in glutamatergic neurons of the aINS, or in the idiopathic polygenic model of autistic behavior BTBR, significantly affects reward-driven responding, impulsivity, or motivational drive under the experimental conditions tested.

### 3.4. *Shank3* deletion alters activity dynamics of glutamatergic neurons in the anterior insular cortex

To determine whether selective deletion of *Shank3* in glutamatergic neurons of the aINS alters neuronal activity, *in vivo* calcium imaging was performed using a miniscope-based recording approach. Neuronal calcium transients were recorded in freely moving mice, in the home cage, and analyzed relative to locomotion states identified using an HMM– based behavioral classification.

Analysis of neuronal activity revealed genotype-dependent differences in calcium signaling between Glu-Shank3-KO-aINS and Control-WT across the three locomotion states of active, explore, and rest (Fig. 5A-C). Specifically, a statistically significant decrease in mean ΔF/F_0_ fluorescence amplitudes in the Glu-Shank3-KO-aINS in comparison to the control was observed across the 3 locomotion states (p<0.05, p<0.01, Fig. 5A-C). In addition, the area under the curve (AUC) of calcium transients (Fig. 5D-F) and the total number of calcium events (Fig. 5J-L) were significantly lower in Glu-Shank3-KO-aINS only in active (p<0.001) and resting states (p<0.001), in comparison to control. No difference was observed in calcium event rate across the 3 locomotion states (Fig. 5G-I).

**Figure 5.**
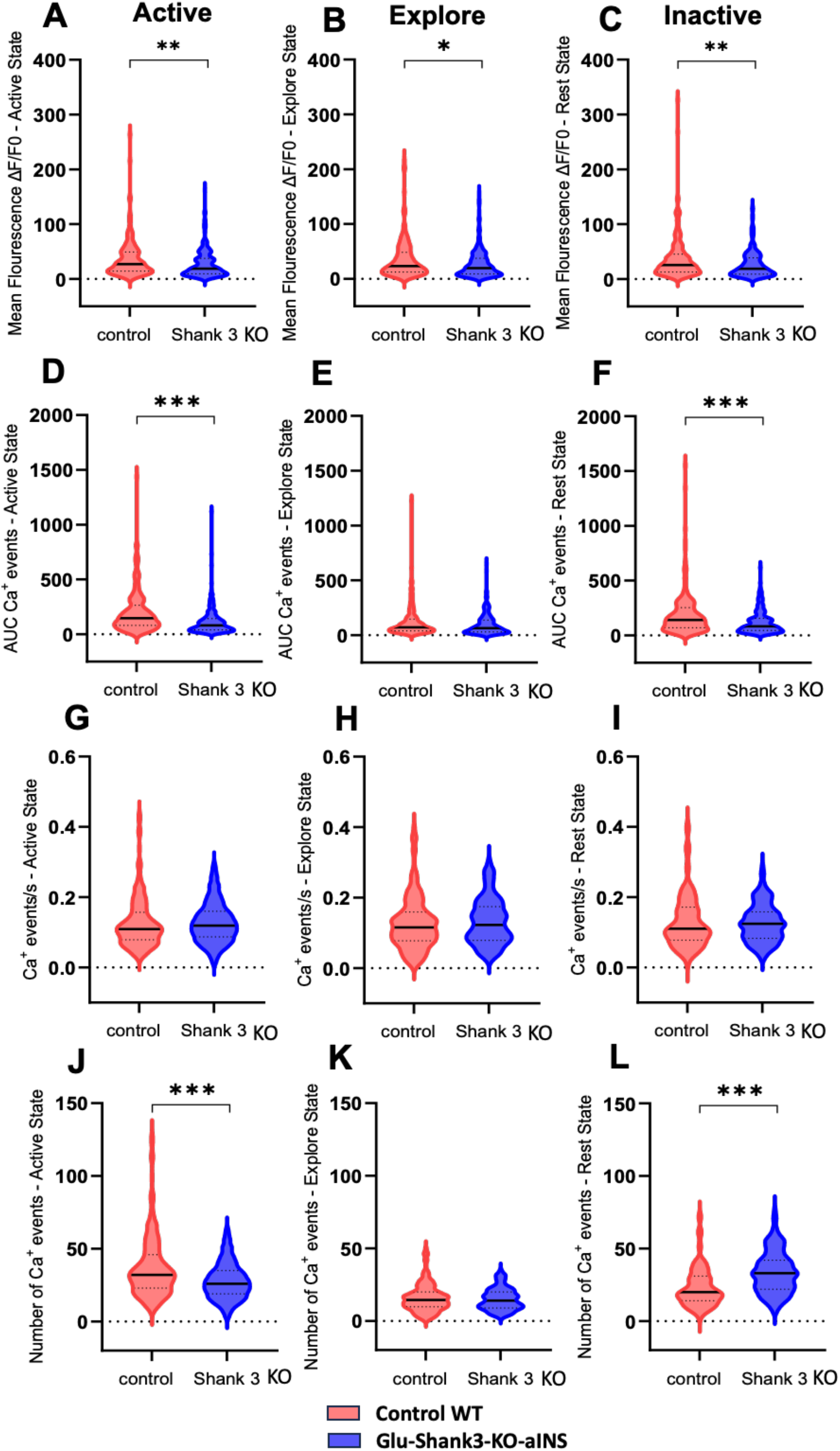
Home-cage neuronal calcium dynamics in anterior insular cortex (aINs) in Control and *Shank3* deletion in glutamatergic neurons across locomotion states. Neuronal calcium activity was recorded in freely moving mice and analyzed across 3 locomotion states: active, explore, and rest. (A–C) Mean ΔF/F_0_ fluorescence amplitude for each locomotion state. (D–F) Area under the curve (AUC) of calcium transients across locomotion states. (G–I) Calcium event rate (events/s) for each locomotion state. (J–L) Total number of calcium events detected in each locomotion state. Each dot represents one neuron. Control: n = 170 neurons; Glu-Shank3-KO-aINs: n = 203 neurons. Data are presented as violin plots with median values. Statistical comparisons were performed using Welch’s t-test (*p < 0.05, **p < 0.01, ***p < 0.001).

Event-based analyses further revealed genotype-dependent differences in neuronal activity patterns (Fig. 6). Classification of neuronal responses based on the association between calcium events and locomotion states showed a reduction in the proportion of neurons categorized as active during the active locomotion state in mutants (Chi-square test, p < 0.01; Fig. 6A, D). In contrast, the proportion of inactive neurons remained similar across the other two behavioral states (Fig. 6A–D). Notably, mutants exhibited a higher proportion of unresponsive neurons during the active locomotion state (p < 0.001). During the exploratory state, mutants displayed a reduced proportion of inactive neurons (p < 0.05), whereas the percentages of active and unresponsive neurons remained comparable to those in controls (Fig. 6B, E). Finally, in the resting state, mutants showed a decreased proportion of active neurons (p < 0.01; Fig. 6C, F), with similar levels of inactive neurons and a significant increase in unresponsive neurons (p < 0.001).

**Figure 6.**
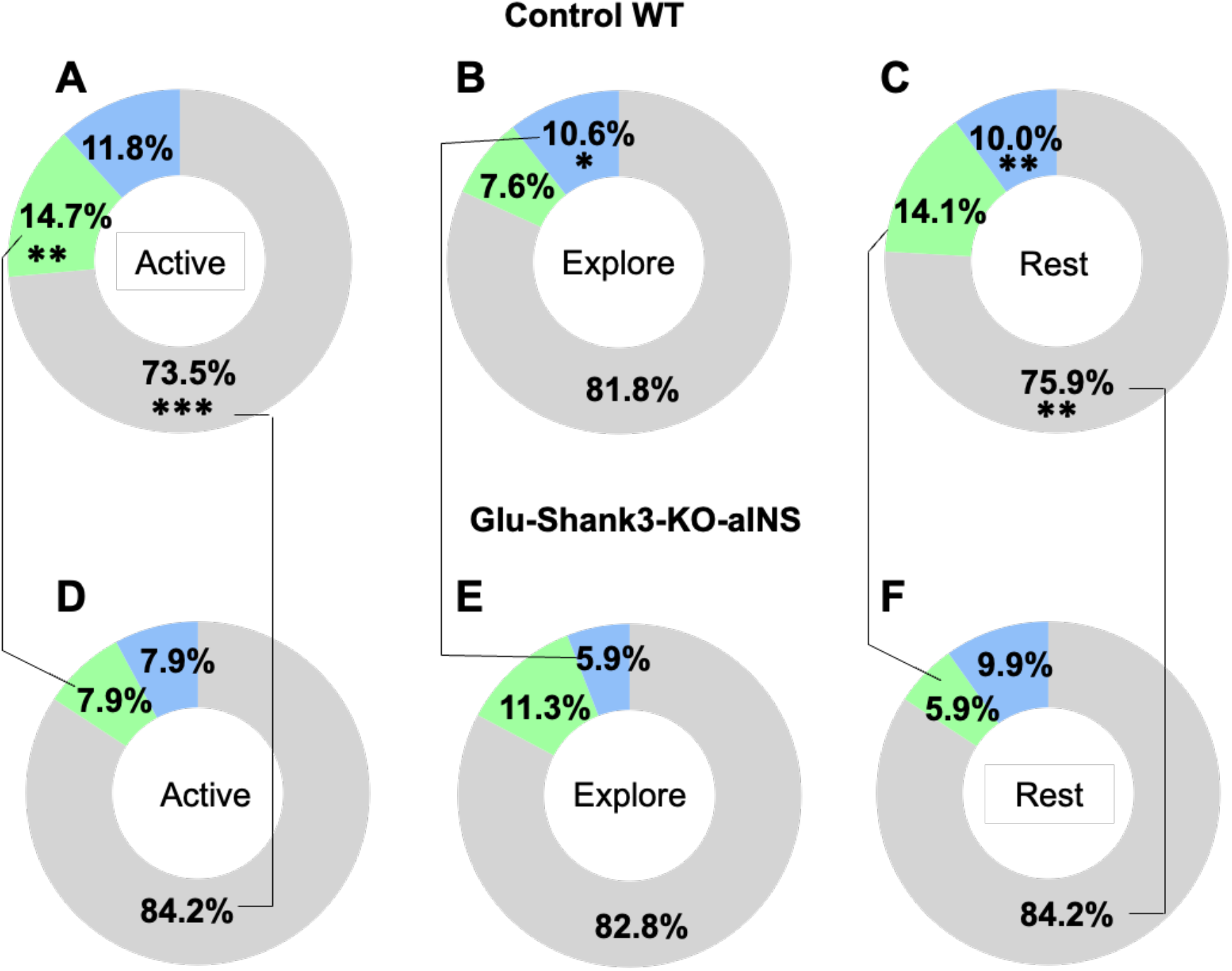
Distribution of neuronal response profiles across locomotion states in Control and *Shank3* deletion mice. Pie charts showing the percentage distribution of neurons classified according to their activity patterns across 3 locomotion states (active, explore, and rest) in their home cage. Neurons were categorized as active (blue), inactive (green), or unresponsive (gray) based on the association between calcium events and locomotion states. (A–C) Distribution of neuronal response types in Control mice (n = 170 neurons) across the 3 locomotion states. (D–F) Distribution of neuronal response types in Shank3KO mice (n = 203 neurons) across the same locomotion states. Data are presented as a pie chart with percentages. Statistical comparisons were performed using the Chi-square test (*p < 0.05, **p < 0.01, ***p < 0.001).

Together, these findings indicate that *Shank3* deletion in glutamatergic neurons of the aINS disrupts neuronal activity dynamics, suggesting dysfunctional organization of insular networks.

## 4. Discussion

In the present study, we demonstrate that selective deletion of *Shank3* in glutamatergic neurons of the aINS is sufficient to induce behavioral alterations relevant to ASD. This brain-specific manipulation produced increased anxiety-like behavior, altered social memory, enhanced repetitive behavior, decreased impulsivity and reward-driven behavior, while leaving social interaction, locomotor activity, general cognitive performance, and motivation largely unaffected. These findings identify glutamatergic neurons of the aINs as a critical circuit locus through which Shank3 dysfunction can disrupt socio-emotional and repetitive behavior.

The aINS plays a central role in integrating interoceptive signals with emotional and motivational states^7^. Through its extensive connectivity with limbic structures, such as the anterior cingulate cortex, amygdala, and striatum, the aINS contributes to the regulation of anxiety, social cognition, and decision-making^5^. Alterations in insular connectivity and activity have been consistently reported in individuals with ASD, suggesting that dysfunction within this region may represent a convergent node in the neural circuitry underlying the disorder^6^. Our findings extend this framework by demonstrating that disruption of a single ASD-associated gene within excitatory neurons in the aINs is sufficient to reproduce several ASD-relevant core symptoms^22^.

Indeed, the behavioral phenotype observed following *Shank3* deletion highlights the importance of glutamatergic synaptic scaffolding in insular circuit function. Shank3 is a key component of the postsynaptic density at excitatory synapses, where it organizes large protein complexes that regulate receptor trafficking, dendritic spine morphology, and synaptic signaling. Loss of Shank3 function disrupts excitatory synaptic transmission and alters neuronal activity patterns in multiple brain regions. By restricting *Shank3* deletion to glutamatergic neurons of the anterior insula, our results indicate that excitatory synaptic scaffold disruption within this specific brain region is sufficient to alter social cognition, anxiety-related, and repetitive behaviors, in addition to reinforcement and impulsivity-like behavior.

Interestingly, the behavioral alterations induced by insular *Shank3* deletion were selective. While anxiety-like behavior, repetitive behavior, and social memory were impaired, sociability, locomotion, and recognition memory remained largely intact. This dissociation suggests that distinct behavioral domains depend on partially segregated circuit mechanisms within the insular network^9^. In particular, the impairment in social novelty recognition observed in our model indicates that Shank3-dependent signaling within aINS excitatory neurons contributes specifically to higher-order aspects of social cognition rather than to basic social approach behavior. In addition, reinforcement and impulsivity-like behaviors were reduced in mutants, suggesting that Shank3-dependent alterations in excitatory neurons of the aINS play a key role in reward processing and impulse control.

Previous studies using total Shank3 knockout models have reported robust alterations in social behavior, repetitive behaviors, and synaptic function^19,23^. In contrast, our aINs-specific deletion of Shank3 was restricted to glutamatergic neurons and resulted in a more selective phenotype, primarily affecting social memory, anxiety-like behavior, reinforcement, impulsivity, and repetitive responses. This difference highlights the importance of circuit-specific contributions to ASD-related behaviors and suggests that distinct brain regions may differentially mediate specific behavioral domains.

Comparison with the idiopathic polygenic ASD BTBR model further highlights the specificity of the brain region manipulation used here. While BTBR mice displayed increased anxiety-like behavior and reduced reinforcement and impulsivity, they did not exhibit robust alterations across all behavioral domains examined in this study. This partial overlap suggests that the insular glutamatergic manipulation captures a specific subset of ASD-related phenotypes rather than broadly reproducing the full behavioral spectrum observed in polygenic idiopathic models. Such differences are expected given the complex genetic and circuit heterogeneity of ASD.

Calcium imaging results provide direct evidence that Shank3 deletion disrupts the functional organization of glutamatergic neurons in the aINs. The overall reduction in calcium signal amplitude, AUC, and number of events, particularly during active and resting states, suggests decreased neuronal engagement rather than a change in firing frequency per se, as event rates remained unchanged. This interpretation is further supported by shifts in neuronal response profiles, characterized by a reduced proportion of active neurons and an increased proportion of unresponsive cells, particularly during behaviorally relevant states. Together, these findings indicate that Shank3 deficiency leads to functional hypoactivity and reduced recruitment of insular glutamatergic neurons, which may underlie the observed behavioral deficits and reflect a disruption in the dynamic encoding of behavioral states within this brain region.

Taken together, our results identify the aINS as a critical circuit node through which Shank3 dysfunction can disrupt several ASD-like core symptoms. By combining brain-type-specific genetic manipulation with behavioral analysis, this study provides direct evidence linking synaptic scaffold disruption in insular excitatory neurons to ASD-relevant phenotypes. These findings advance our understanding of how genetic risk factors for autism translate into circuit-level dysfunction and highlight the aINS Sahnk3 as a key target for future investigations into the neural mechanisms underlying ASD.

## Acknowledgements

This work was supported by CRC1193 “Neurobiology of resilience” to B.L.; ERA-NET Neuron/Bundesministerium für Bildung und Forschung (BMBF)/DLR Projektträger ref. 01EW2203A to B.L. and ref. 01EW2203B to M.J.S. Spanish “Ministerio de Ciencia, Innovación y Universidades, Agencia Estatal de Investigación (AEI)” (PID2020-120029GB-I00/MICIN/AEI/10.13039/501100011033, RD21/0009/0019 and PDI2023-1511680B-C21), the Spanish “Instituto de Salud Carlos III, RETICS-RTA” (#RD12/0028/0023), the “Generalitat de Catalunya, AGAUR” (#2020 SGR), “ICREA-Acadèmia” (#2025) and the Spanish “Ministerio de Sanidad, Servicios Sociales e Igualdad”, “Plan Nacional Sobre Drogas of the Spanish Ministry of Health” (#PNSD-2022) to R.M., “la Caixa Health” LCR/PR/HR22/5240017 to R.M. and E.M-G., “Plan Nacional Sobre Drogas of the Spanish Ministry of Health” (#PNSD-2019I006, #PNSD-2023I040) and Spanish “Ministerio de Ciencia e Innovación” (ERA-NET) PCI2021-122073-2A to E.M-G.

We are very grateful to R. Martín and F. Porrón for their technical support. Figures 1 and 4 were created with BioRender.

## Author contributions

E. M.-G. and R. M. conceived and designed the behavioral studies with input from B. L. and I. R. A.; B. L. generated the conditional transgenic mice; P. M.-A. and G. H. performed the behavioral experiments and the statistical analyses and graphs with the supervision of E.M.-G., R.M. and M. J. S.; I. R. A performed the immunofluorescence studies; I. M.-B. performed the calcium imaging experiments in collaboration with Z. M. and T. G and with the supervision of R. A. Bioinformatic analyses were performed by J. P.-V; E. M.-G., P. M.-A., G. H. and R. M. wrote the manuscript and M. S., I.R.A and B. L. provided critical review of the manuscript with inputs from all the other authors.

## Data availability

Individual data points are graphed in the main figures. All the relevant data that support this study are available from the corresponding author to any interested researcher upon reasonable request.

## Figures

**Supplementary Figure 1.**
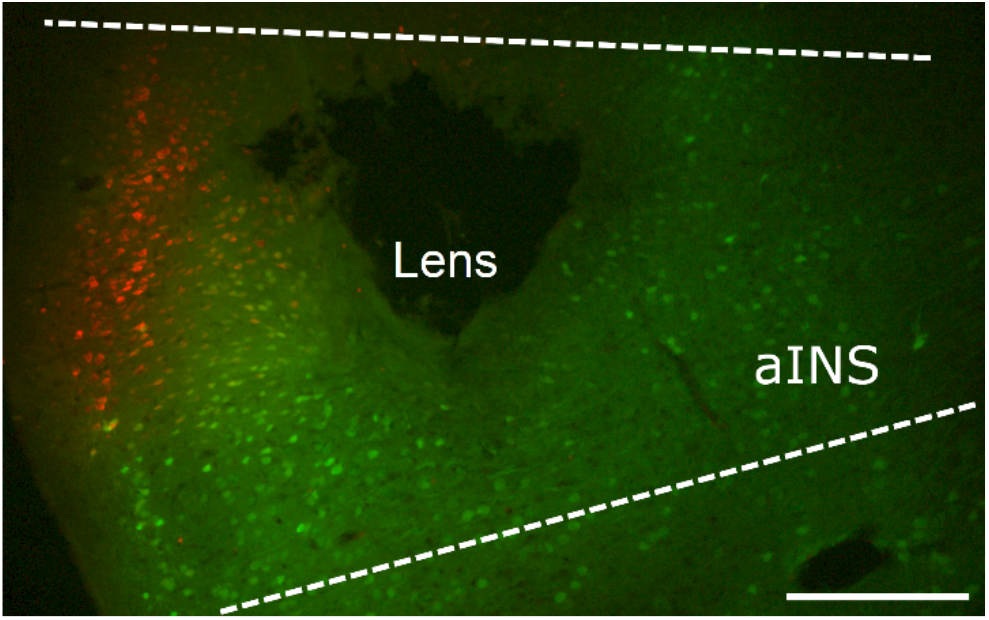
Representative fluorescent image of Cre- and GCaMP6f-expressing neurons in the aINS and the lens implantation site. Scale bar: 300 µm.

## References

1 Lord, C. et al. Autism spectrum disorder. Nat Rev Dis Primers 6, 5 (2020). 10.1038/s41572-019-0138-4

2 Hyman, S. L., Levy, S. E., Myers, S. M., Council On Children With Disabilities, S. O. D. & Behavioral, P. Identification, Evaluation, and Management of Children With Autism Spectrum Disorder. Pediatrics 145 (2020). 10.1542/peds.2019-3447

3 Nardi, L. et al. Neuroanatomical changes of ionotropic glutamatergic and GABAergic receptor densities in male mice modeling idiopathic and syndromic autism spectrum disorder. Front Psychiatry 14, 1199097 (2023). 10.3389/fpsyt.2023.1199097

4 Reim, D. et al. Proteomic Analysis of Post-synaptic Density Fractions from Shank3 Mutant Mice Reveals Brain Region Specific Changes Relevant to Autism Spectrum Disorder. Front Mol Neurosci 10, 26 (2017). 10.3389/fnmol.2017.00026

5 Gogolla, N. The insular cortex. Curr Biol 27, R580–R586 (2017). 10.1016/j.cub.2017.05.010

6 Gehrlach, D. A. et al. A whole-brain connectivity map of mouse insular cortex. Elife 9 (2020). 10.7554/eLife.55585

7 Namkung, H., Kim, S. H. & Sawa, A. The Insula: An Underestimated Brain Area in Clinical Neuroscience, Psychiatry, and Neurology. Trends Neurosci 40, 200–207 (2017). 10.1016/j.tins.2017.02.002

8 Di Martino, A. et al. The autism brain imaging data exchange: towards a large-scale evaluation of the intrinsic brain architecture in autism. Mol Psychiatry 19, 659–667 (2014). 10.1038/mp.2013.78

9 Uddin, L. Q. & Menon, V. The anterior insula in autism: under-connected and under-examined. Neurosci Biobehav Rev 33, 1198–1203 (2009). 10.1016/j.neubiorev.2009.06.002

10 Cochoy, D. M. et al. Phenotypic and functional analysis of SHANK3 stop mutations identified in individuals with ASD and/or ID. Mol Autism 6, 23 (2015). 10.1186/s13229-015-0020-5

11 Monteiro, P. & Feng, G. SHANK proteins: roles at the synapse and in autism spectrum disorder. Nat Rev Neurosci 18, 147–157 (2017). 10.1038/nrn.2016.183

12 Leblond, C. S. et al. Meta-analysis of SHANK Mutations in Autism Spectrum Disorders: a gradient of severity in cognitive impairments. PLoS Genet 10, e1004580 (2014). 10.1371/journal.pgen.1004580

13 Phelan, K. & McDermid, H. E. The 22q13.3 Deletion Syndrome (Phelan-McDermid Syndrome). Mol Syndromol 2, 186–201 (2012). 10.1159/000334260

14 Sheng, M. & Kim, E. The postsynaptic organization of synapses. Cold Spring Harb Perspect Biol 3 (2011). 10.1101/cshperspect.a005678

15 Mei, Y. et al. Adult restoration of Shank3 expression rescues selective autistic-like phenotypes. Nature 530, 481–484 (2016). 10.1038/nature16971

16 Peca, J. et al. Shank3 mutant mice display autistic-like behaviours and striatal dysfunction. Nature 472, 437–442 (2011). 10.1038/nature09965

17 Wang, X. et al. Synaptic dysfunction and abnormal behaviors in mice lacking major isoforms of Shank3. Hum Mol Genet 20, 3093–3108 (2011). 10.1093/hmg/ddr212

18 Meyza, K. Z. & Blanchard, D. C. The BTBR mouse model of idiopathic autism - Current view on mechanisms. Neurosci Biobehav Rev 76, 99–110 (2017). 10.1016/j.neubiorev.2016.12.037

19 Drapeau, E., Riad, M., Kajiwara, Y. & Buxbaum, J. D. Behavioral Phenotyping of an Improved Mouse Model of Phelan-McDermid Syndrome with a Complete Deletion of the Shank3 Gene. eNeuro 5 (2018). 10.1523/ENEURO.0046-18.2018

20 Domingo-Rodriguez, L. et al. A specific prelimbic-nucleus accumbens pathway controls resilience versus vulnerability to food addiction. Nat Commun 11, 782 (2020). 10.1038/s41467-020-14458-y

21 Martin-Garcia, E., Domingo-Rodriguez, L. & Maldonado, R. An Operant Conditioning Model Combined with a Chemogenetic Approach to Study the Neurobiology of Food Addiction in Mice. Bio Protoc 10, e3777 (2020). 10.21769/BioProtoc.3777

22 Vicidomini, C. et al. Pharmacological enhancement of mGlu5 receptors rescues behavioral deficits in SHANK3 knock-out mice. Mol Psychiatry 22, 784 (2017). 10.1038/mp.2016.70

23 Betancur, C. & Buxbaum, J. D. SHANK3 haploinsufficiency: a “common” but underdiagnosed highly penetrant monogenic cause of autism spectrum disorders. Mol Autism 4, 17 (2013). 10.1186/2040-2392-4-17

